## Supplemental Materials for "Directed evolution of the rRNA methylating enzyme Cfr reveals molecular basis of antibiotic resistance"

***Short title:*** **Directed evolution of the Cfr resistance enzyme**

Kaitlyn Tsai, Vanja Stojković, Lianet Noda-Garcia, Iris D. Young, Alexander G. Myasnikov, Jordan Kleinman, Ali Palla, Stephen N. Floor, Adam Frost, James S. Fraser, Dan S. Tawfik, Danica Galonić Fujimori

**This file includes:**

A. Supplementary Figures 1-12
B. Supplementary Tables 1-6
C. References for Supplementary Materials

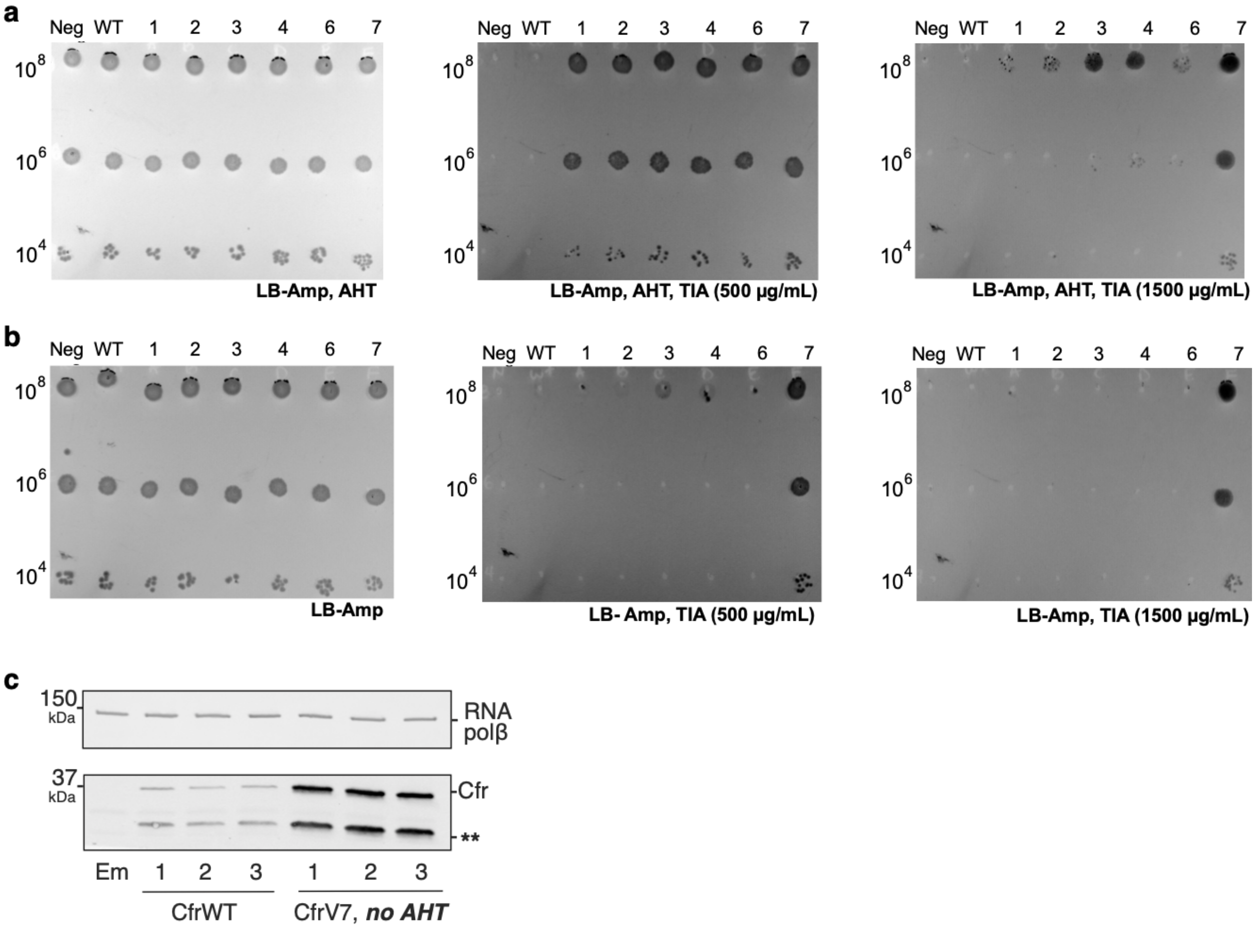


**Supplementary Fig. 1. CfrV7 does not require an inducer for resistance or expression.** (**a**) Antibiotic susceptibility testing of *E. coli* transformed with pZA plasmid encoding Cfr variants. Dose-dependent susceptibility testing toward tiamulin (TIA) with plates containing ampicillin (Amp, 100 µg/mL) and anhydrotetracycline (AHT, 20 ng/mL) to induce Cfr expression. (**b**) Dose-dependent susceptibility testing towards TIA with plates containing Amp but lacking AHT. Neg designates empty pZA plasmid. WT and 1-7 designates pZA encoding CfrWT-His$_6$ or Cfr variants 1-7. For panels (a) and (b), results are representative of three biological replicates. (**c**) Protein expression of full-length CfrV7 without AHT inducer compared to CfrWT with AHT inducer detected by immunoblotting against a C-terminal FLAG tag. Em = empty vector control.

a

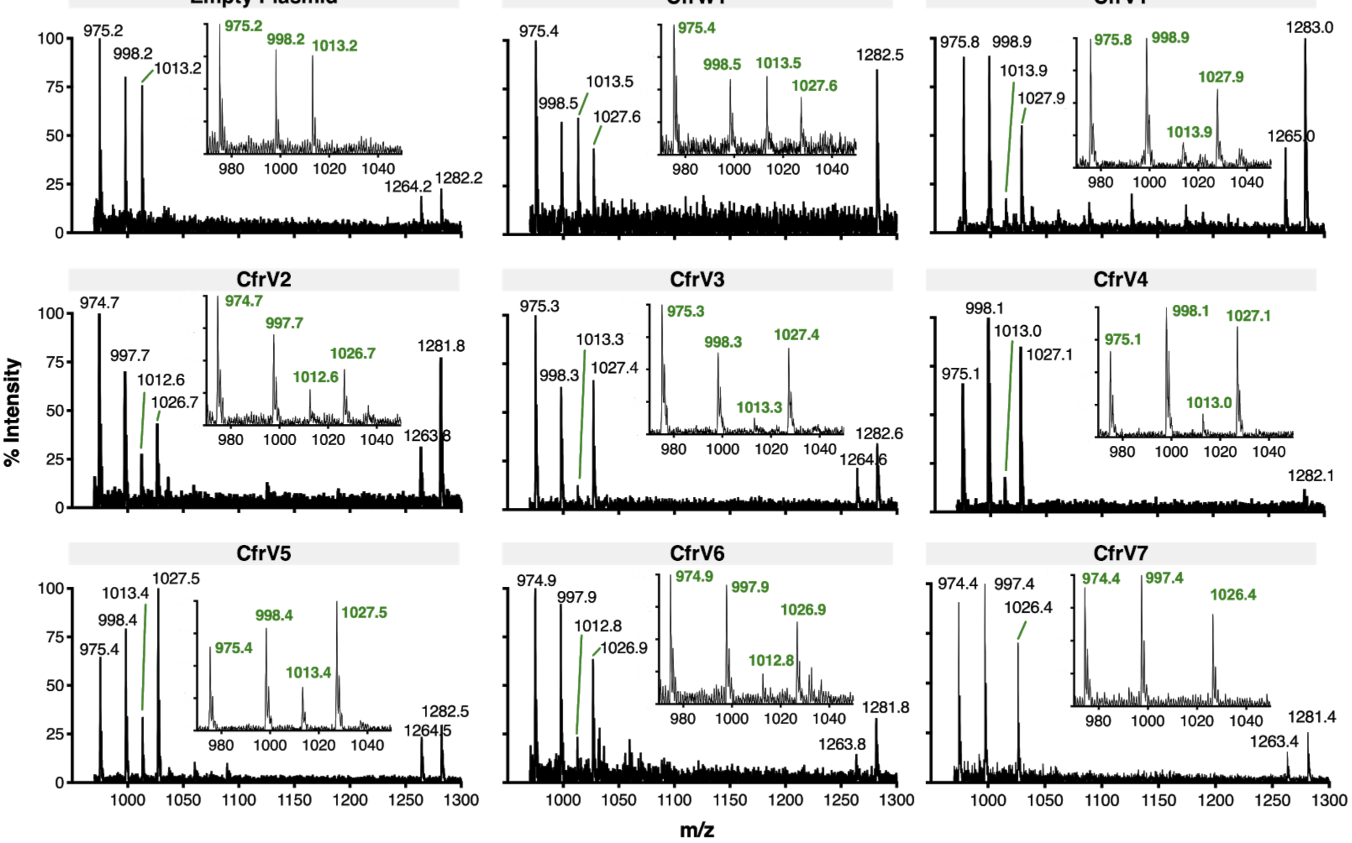


b

| 23S rRNA Fragment | Sequence | m/z |
|---|---|---|
| 2482-2484 | ACG | 998.16 |
| 2491-2494 | UUUG | 1282.14 |
| 2496-2502 | CACmCUCG | 2233.32 |
| **2503-2505** | **m²AΨG** | **1013.16** |
| | **m²m⁸AΨG** | **1027.18 (+14)** |
| 2506-2508 | UCG | 975.13 |
| 2510-2520 | CUCAUCACAUC | 3450.53 |

**Supplementary Fig. 2. MALDI-TOF mass spectra of 23S rRNA fragments produced by oligo-protection and RNase $T_1$ digestion.** (**a**) RNA isolated from *E. coli* with empty pZA vector or expressing CfrWT and evolved Cfr variants CfrV1-V7. Spectra are representative of 2-3 biological replicates. Endogenously modified ($m^2$A) and Cfr-modified rRNA fragments ($m^2m^8$A) correspond to m/z values of 1013 and 1027, respectively. Spectra are representative of 2-3 biological replicates. (**b**) Expected rRNA fragments and m/z values produced by RNase $T_1$ digestion of C2480–C2520 23S rRNA nucleotides after oligo-protection. Peaks ~1264 m/z corresponds to the cyclic phosphate of the UUUG rRNA fragment. Cm is methylated cytosine, Ψ is pseudouridine, $m^2$A is 2-methyladenosine, $m^2m^8$A is 2,8-dimethyladenosine.

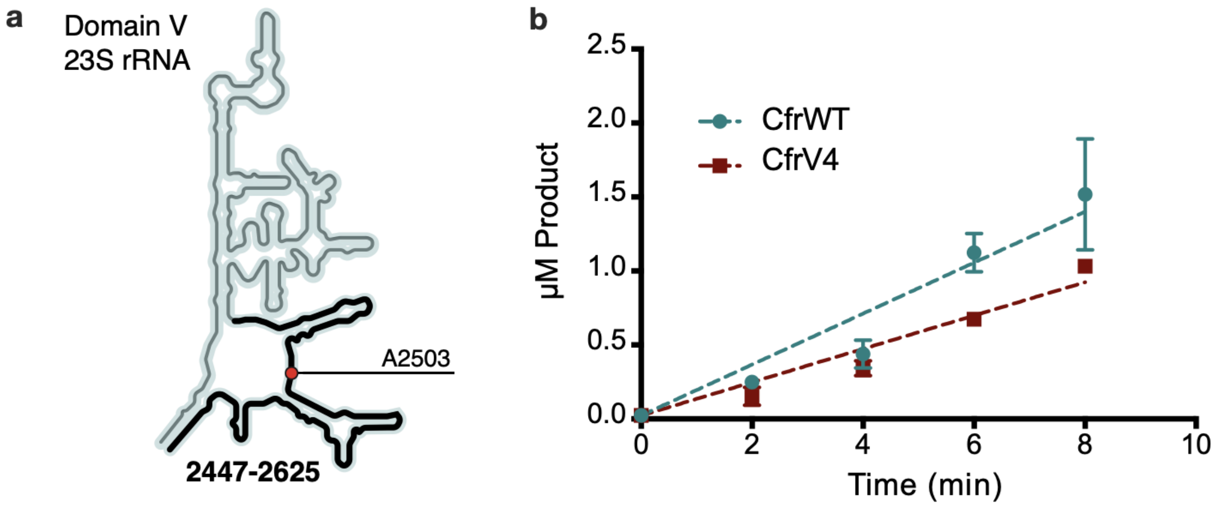


**Supplementary Fig. 3. *In vitro* characterization of CfrWT and CfrV4.** (**a**) Schematic representation of 23S rRNA domain V secondary structure highlighting the rRNA fragment 2447-2625 used in the kinetic assay. Position of the Cfr substrate nucleotide A2503 is indicated by a red circle. (**b**) Time-dependent formation of methylated rRNA product by CfrWT or evolved CfrV4 in reactions containing 5 µM apo-reconstituted Cfr enzyme, 100 µM 23S rRNA fragment 2447-2625, and 2 mM [$^3$H-methyl] *S*-adenosylmethionine. Values are presented as the average and standard error of two replicates. Turnover numbers for CfrWT and CfrV4 are 3.45 x $10^{-2}$ ± 3.2 x $10^{-3}$ $min^{-1}$ and 2.25 x $10^{-2}$ ± 1.3 x $10^{-3}$ $min^{-1}$, respectively.

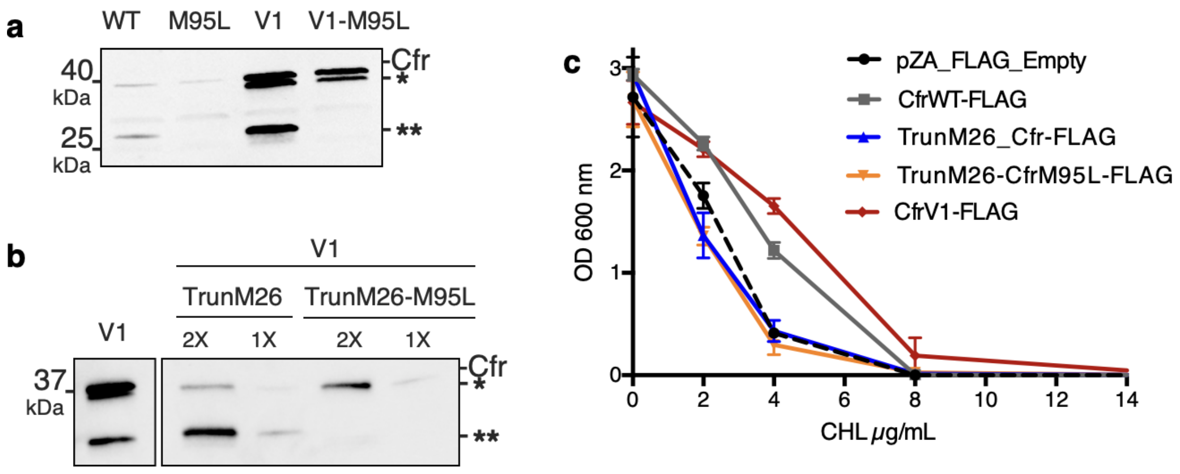


**Supplementary Fig. 4. N-terminally truncated Cfr products arise from internal Met translation start sites and do not contribute to resistance**. (**a**) Detection of Cfr protein products by immunoblotting against C-terminal FLAG tag. Directed evolution mutation I26M is sufficient to produce the higher-molecular weight truncated species (denoted by *). Mutation of M95L (AUG -> CUG) abolishes production of smaller truncated species (denoted by **). Blot is representative of two biological replicates. (**b**) Detection of Cfr protein products by immunoblotting against C-terminal FLAG tag with ANTI-FLAG M2-Peroxidase antibody (Sigma). TrunM26 refers to a Cfr construct where amino acids 1-25 have been eliminated. This construct yields predominately species **, with * as the minor expressed protein. TrunM26-M95L eliminates amino acids 1-25 and also removes internal Met start site through M95L mutations, yielding species *. (**c**) Dose-dependent growth inhibition of *E. coli* expressing N-terminally truncated Cfrs in the presence of chloramphenicol (CHL). Results presented as the average of two biological replicates with standard error.

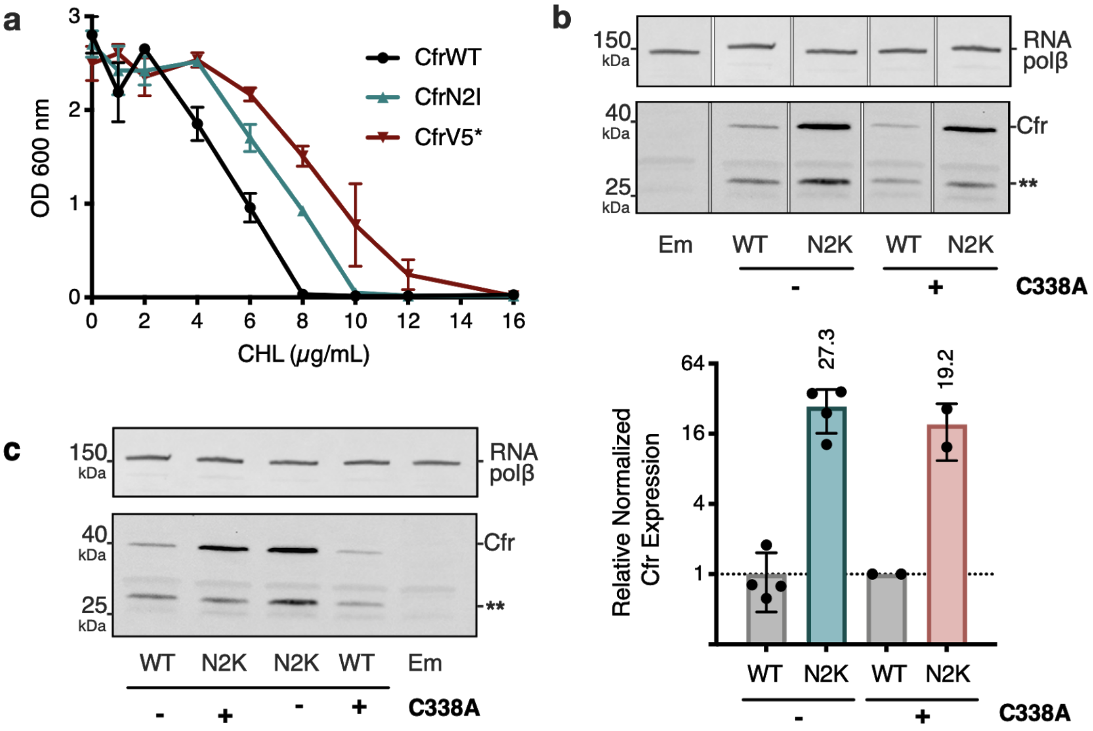


**Supplementary Fig. 5. Investigations into second position Cfr mutations N2I and N2K.** (**a**) Dose-dependent growth inhibition of *E. coli* expressing CfrN2I compared to CfrV5 towards chloramphenicol (CHL). Results presented as an average of three biological replicates with standard error. (**b**) Relative expression of CfrN2K compared to CfrWT, in the presence or absence of an additional C338A mutation that renders Cfr catalytically inactive. Therefore, in constructs carrying the C338A mutation, Cfr constructs are produced by ribosomes that lack the $m^8$A2503 modification. Signal of full-length Cfr protein was normalized to housekeeping protein RNA polymerase β-subunit, presenting the average of two biological replicates and standard deviation on a $\log_2$ axis. Asterisks denote truncated Cfr products that do not contribute to resistance and were not included in quantification. (**c**) Original, uncropped image of panel (a). Em = empty vector control.

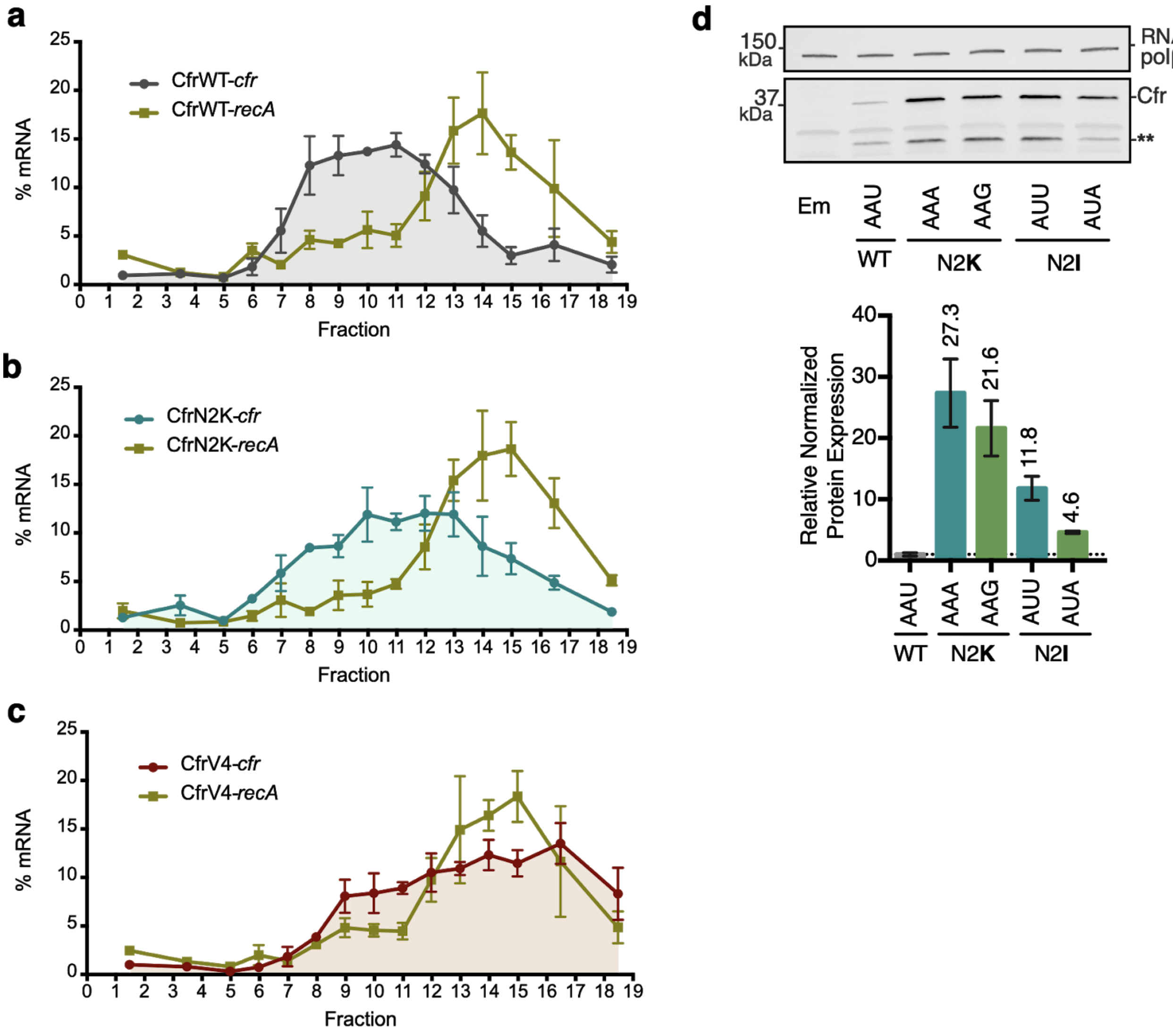


**Supplementary Fig. 6. Investigating translation of Cfr mutants**. Distribution of Cfr and recA transcripts across polysome profiles from *E. coli* expressing (**a**) pZA-encoded CfrWT, (**b**) CfrN2K and (**c**) CfrV4. Transcript levels for each fraction isolated from a 10-55% sucrose gradient were determined by RT-qPCR and normalized by a luciferase mRNA control spike-in. Values are presented as the average of three biological replicates with standard error. (**d**) Relative protein expression of Cfr with second codon mutations compared to CfrWT. Signal corresponding to full-length Cfr protein was normalized to housekeeping protein RNA polymerase β-subunit and averaged from three biological replicates with standard error. Asterisks denote the truncated Cfr products that do not contribute to resistance and were not included in quantification. Em = empty vector control.

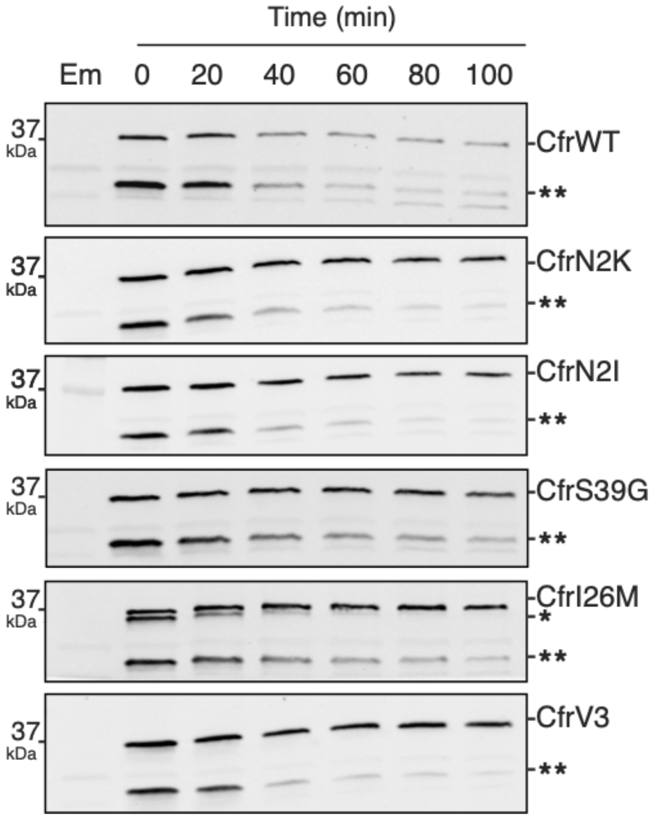


**Supplementary Fig. 7. Degradation of Cfr protein products.** Protein degradation kinetics of CfrWT, single mutations CfrN2K/N2I/S39G/I26M, and evolved variant CfrV3 in *E. coli* after halting expression by rifampicin and immunoblotting against the C-terminal FLAG tag. Presented western blots are expanded images of those presented in Fig. 4 to display degradation of the Cfr truncation denoted by two asterisks (**). Em = empty vector control.

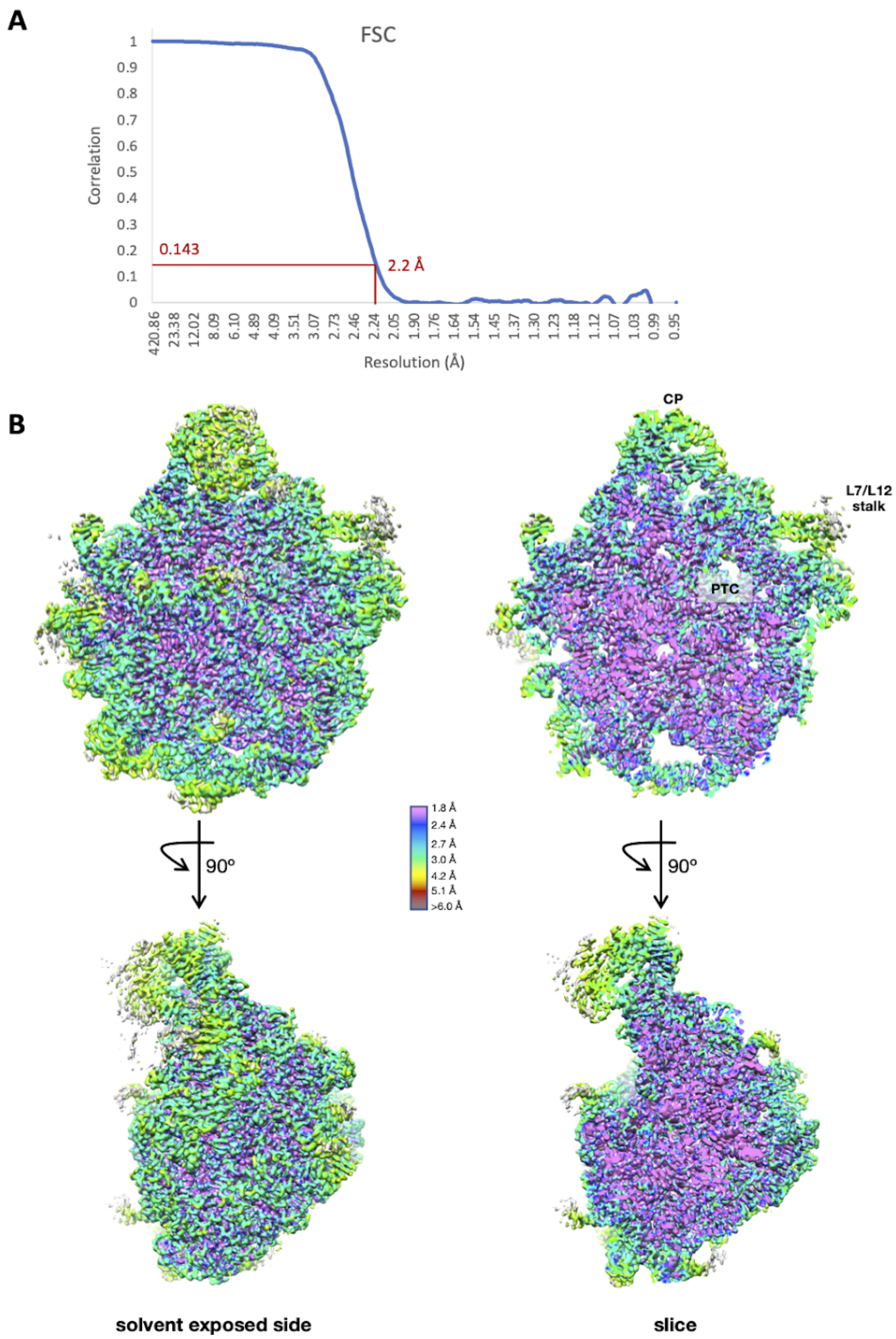


**Supplementary Fig. 8. Cryo-EM data collection and processing of the Cfr-modified ribosome.** (**a**) Fourier shell correlation (FSC) curve indicating the overall resolution of the *E. coli* 50S subunit using the FSC = 0.143 gold standard. (**b**) 3D cryo-EM map of the *E. coli* 50S subunit colored according to local resolution, highlighting the L7/L12 stalk and peptidyl transferase center (PTC). Local resolution map was calculated with ResMap. Figure was prepared using UCSF Chimera 1.13 (Pettersen et al., 2004).

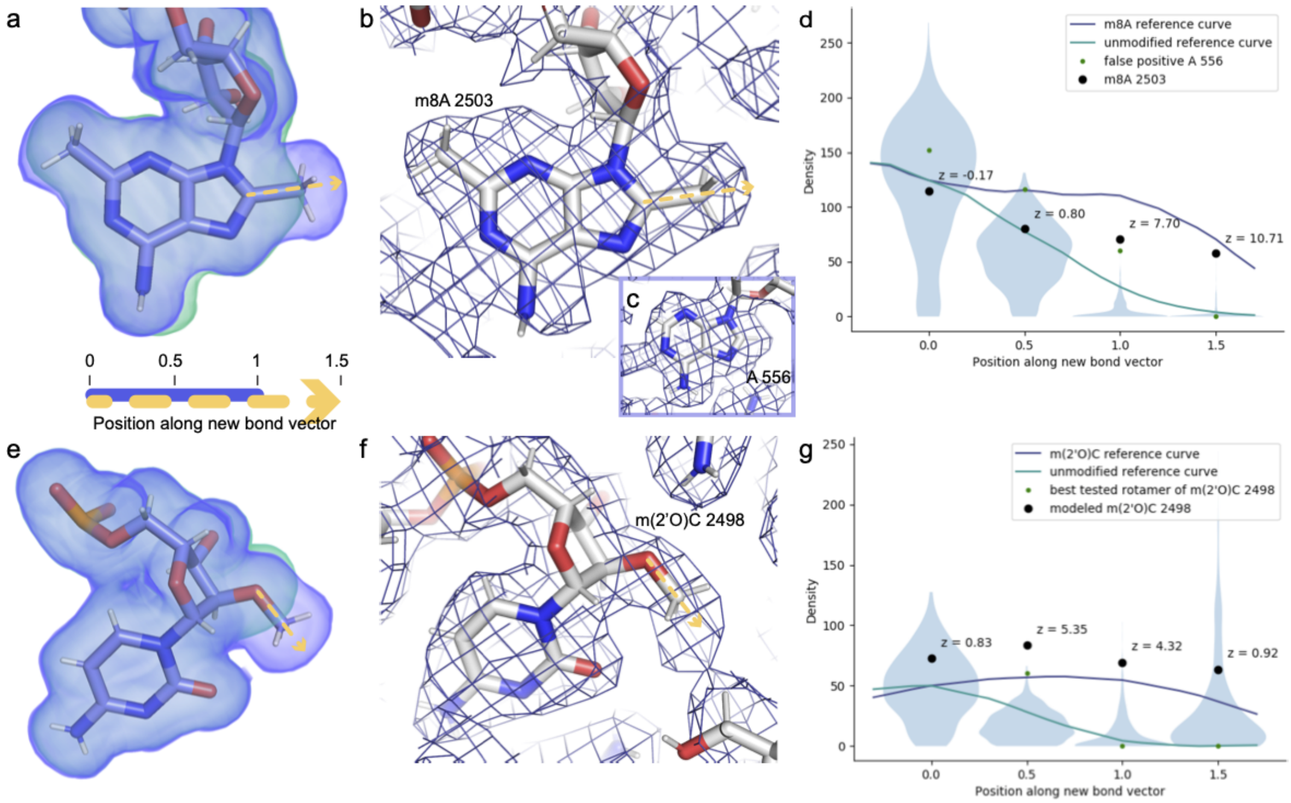


**Supplementary Fig. 9. Cross-validation of methylations on C8 of A2503 and 2′O of C2498 from the cryoEM density map.** Ideal, noise-free densities were calculated for the post-transcriptionally modified (purple) and unmodified (green) nucleotides (**a, e**). We can distinguish which of these maps better matches the experimental map by examining the dropoff in density when moving from the reference atom (C8 or 2′O) toward and beyond the methyl group. To this end, the noise-free calculated densities were used to generate reference curves for these two modifications (**d, g**). These curves were scaled to match the mean density at the reference atom for all instances of adenosines (**d**) or cytosines (**g**) in the experimental structure, and for those same nucleotides, densities at four selected positions along the vector are also shown in violin plots (**d, g**). Based on these four densities as well as calculated difference density at the 1.0 position and the correlation coefficient between the calculated and experimental map for the entire nucleotide, the program qPTxM for detection of posttranscriptional modifications identified two adenosines where the map supported C8 methylation: A2503 and a false positive A556 (**b-d**). Investigation of the densities at these two sites confirmed that the shape of the density dropoff curve for A2503 more closely matched the methylated reference curve while that of A556 more closely matched the unmethylated reference curve, and that the latter site was identified as a strong candidate primarily due to the strong density at all atoms. qPTxM identified no cytosines where the map supported 2′O methylation (**g**). The densities along the methylation bond vector more closely match the unmodified than the methylated reference curve at C2498 (**f**) when the methyl group is placed at any of the rotameric positions (green dots), but along the modeled O-Me bond (black dots), the dropoff closely matches the shape of the methylated reference curve (**g**). Plots are annotated with Z-scores for the A2503 and C2498 densities relative to all adenosines and cytosines in the map.

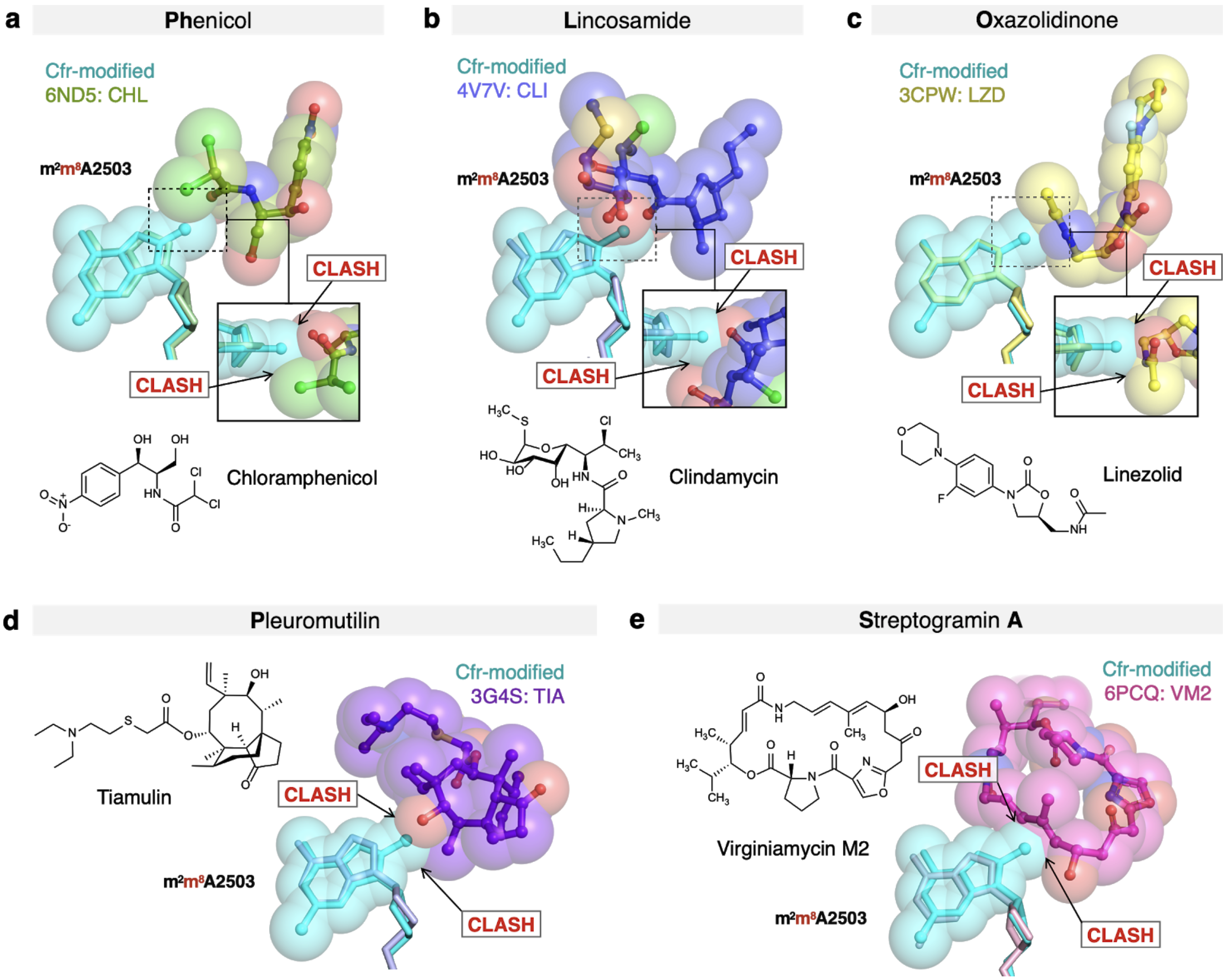


**Supplementary Fig. 10. Molecular basis of Cfr-mediated resistance to PhLOPS$_A$ antibiotics.** Structural overlay of Cfr-modified *E. coli* 50S ribosome (cyan) and ribosomes in complex with (**A**) chloramphenicol (CHL); PDB 6ND5, (**B**) clindamycin (CLI); PDB 4V7V, (**C**) linezolid (LZD); PDB 3CPW, (**D**) tiamulin (TIA) PDB 3G4S, and (**E**) virginiamycin M2 (VM2); PDB 6PCQ. Inserts are close up views of the steric clashes between m$^8$A2503 and corresponding PTC-targeting antibiotics.

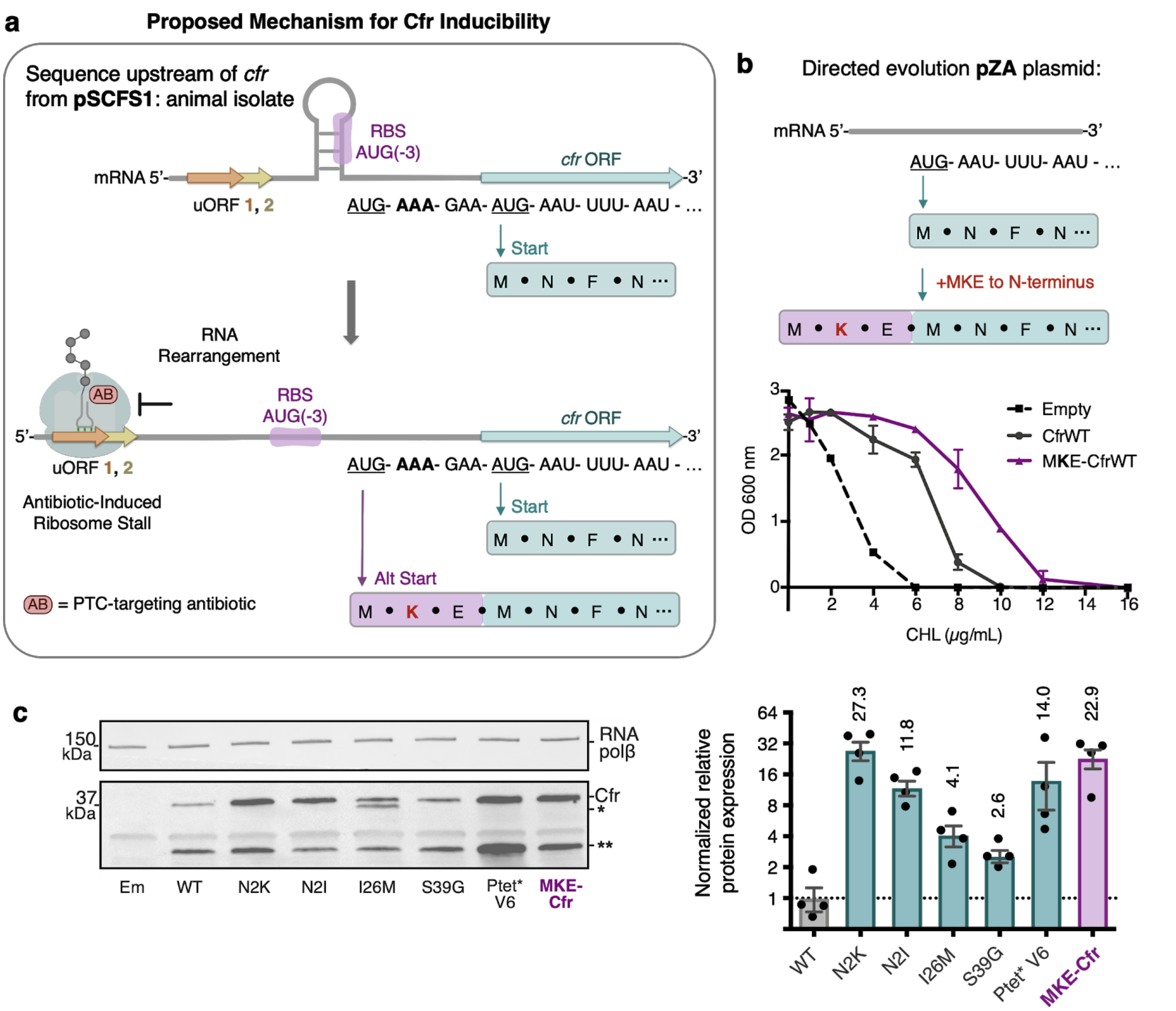


**Supplementary Fig. 11. Start codon selection as a proposed mechanism of Cfr inducibility.** (**a**) Sequence upstream of *cfr* from pSCFS1 (Accession: AJ579365) resistance plasmid from an animal-derived *S. sciuri* isolate. The upstream region contains 2 overlapping upstream ORFs (uORFs) followed by an RNA structural element. Proposed antibiotic-induced ribosome stalling at uORF1/2 and RNA rearrangement could reveal the occluded RBS, allowing translation to initiate at AUG(-3), adding an MKE polypeptide to the N-terminus of Cfr. (**b**) Addition of an N-terminal MKE peptide to Cfr in the context of the pZA plasmid where expression is controlled by the non-native tetracycline-inducible promoter, $P_{tet}$. Growth inhibition of *E. coli* with pZA-encoded MKE-Cfr in the presence of CHL determined from two biological replicates with standard error. (**c**) Relative protein expression of full-length MKE-Cfr compared to full-length CfrWT detected by immunoblotting against a C-terminal FLAG tag and quantification of top Cfr bands. Signal was normalized to housekeeping protein RNA polymerase β-subunit. Data is presented as the average of four biological replicates with standard deviation on a $\log_2$ axis. Em = empty vector control.

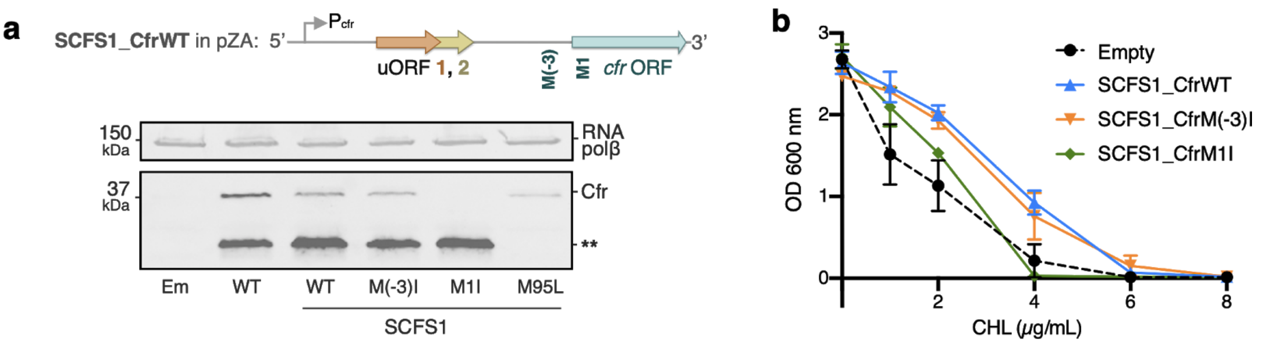


**Supplementary Fig. 12. Investigations into Cfr translation start sites in the pSCFS1 sequence context.** (**a**) Schematic of the SCFS1_CfrWT construct which includes the upstream ORF sequence from pSCFS1 and native $P_{cfr}$ promoter (Accession: AJ579365). Expression of CfrWT from $P_{tet}$ compared to SCFS1_CfrWT and corresponding methionine mutants without inducer. Cfr protein products were detected by immunoblotting against the C-terminal FLAG tag. M(-3)I corresponds to AUG -> AUC mutation of the MKE upstream methionine; M1I corresponds to AUG -> AUC mutation of the annotated start site; M95L corresponds to AUG -> CUG mutation of the M95 internal start site. Mutation of the annotated start site M1I, but not the upstream start site M(-3)I, abolishes detectable expression of full-length Cfr. Asterisks denote the truncated Cfr product from internal translation initiation at M95 that does not contribute to resistance. Presented blot is representative of two biological replicates. (**b**) Dose-dependent growth inhibition of *E. coli* expressing SCFS1_CfrWT and corresponding methionine mutants towards chloramphenicol (CHL) determined from two biological replicates with standard error. Mutation of the annotated start site M1I, but not the upstream start site M(-3)I, impacts antibiotic resistance under these experimental conditions.

| Primer Name | Application | Sequence |
|---|---|---|
| GS1-1XFLAG | Fastcloning Insertion | Fwd: 5’-GATTACAAGGATGACGACGATAAGTGAGCGGCCGCAA ACATGGTAC-3’<br>Rev: 5’-CATCCTTGTAATCGCTACCACCACCTTGGCTATTTTGAT AATTACC-3’ |
| CfrN2K(AAA) | Mutagenesis | Fwd: 5’-AGCTAACCGATGAA**A**TTTAATAATAAAAC-3’<br>Rev: 5’-GTTTTATTATTAAA**T**TTCATCGGTTAGCT-3’ |
| CfrN2K(AAG) | Mutagenesis | Fwd: 5’-AGCTAACCGATGAA**G**TTTAATAATAAAAC-3’<br>Rev: 5’-GTTTTATTATTAAA**C**TTCATCGGTTAGCT-3’ |
| CfrN2I(AUU) | Mutagenesis | Fwd: 5’-AGCTAACCGATGA**TT**TTTAATAATAAAAC-3’<br>Rev: 5’-GTTTTATTATTAAA**AA**TCATCGGTTAGCT-3’ |
| CfrN2I(AUA) | Mutagenesis | Fwd: 5’-AGCTAACCGATGA**TA**TTTAATAATAAAAC-3’<br>Rev: 5’-GTTTTATTATTAAA**TA**TCATCGGTTAGCT-3’ |
| CfrI26M | Mutagenesis | Fwd: 5’-TGAGCCTGATTATAGAAT**G**AAACAAATAACCAATGCG-3’<br>Rev: 5’-CGCATTGGTTATTTGTTT**C**ATTCTATAATCAGGCTCA-3’ |
| CfrS39G | Mutagenesis | Fwd: 5’-GATTTTTAAACAAAGAATT**G**GTCGATTTGAGGATATGAA-3’<br>Rev: 5’-TTCATATCCTCAAATCGAC**C**AATTCTTTGTTTAAAAATC-3’ |
| MK(AAA)E-Cfr | Fastcloning Insertion | Fwd: 5’-**ATGAAAGAA**ATGAATTTTAATAATAAAACAAAGTATG GTAAAATACAG-3’<br>Rev: 5’-ATTCATTTCTTTCATCGGTTAGCTTATCGATAC-3’ |
| MK(AAG)E-Cfr | Fastcloning Insertion | Fwd: 5’-**ATGAAGGAAA**TGAATTTTAATAATAAAACAAAGTATG GTAAAATACAG-3’<br>Rev: 5’-ATTCATTTCCTTCATCGGTTAGCTTATCGATAC-3’ |
| CfrM95L | Mutagenesis | Fwd: 5’-GTAGAAACGGTAAAC**C**TGAAGTATAAAGCAG-3’<br>Rev: 5’-CTGCTTTATACTTCA**G**GTTTACCGTTTCTAC-3’ |
| TrunM26-Cfr | Fastcloning Deletion | Fwd: 5’-TCGATAAGCTAACCGATGAAACAAATAACCAATGCG-3’<br>Rev: 5’-CGGTTAGCTTATCGATACCGTCGACC-3’ |
| CfrC338A | Mutagenesis | Fwd: 5’-_ATTGACGCTGCT**GC**TGGTCAATTATATG-3’<br>Rev: 5’-_CATATAATTGACCA**GC**AGCAGCGTCAAT-3’ |
| cfr | RT-qPCR | Fwd: 5’-AGCAGAGCAAAATTCAGAGCAAGT-3’<br>Rev: 5’-TCCAATGTCGCCTGTAGCACAA-3’<br>Length of amplicon: 169 bp |
| luc<br>Accession no: X65316.2 | RT-qPCR | Fwd: 5’-AGATCGTGGATTACGTCGCC-3’<br>Rev: 5’-TGGACTTTCCGCCCTTCTTG-3’<br>Length of amplicon: 156 bp |
| recA<br>Accession no: CP037857.1 | RT-qPCR | Fwd: 5’-ATCGCCTGGCTCATCATACG-3’<br>Rev: 5’-GCACTGGAAATCTGTGACGC-3’<br>Length of amplicon: 152 bp |
| CfrM(-3)I | Mutagenesis | Fwd: 5’-TTACCACTAGAGCAAATTGTGAAAGGAT**C**AAAGAAATG-3’<br>Rev: 5’-CCTGTATTTTACCATACTTTGTTTTATTATTAAAATTC ATTTCTTT**G**ATCCTTTC-3’ |
| CfrM1I | Mutagenesis | Fwd: 5’-CACTAGAGCAAATTGTGAAAGGATGAAAGAAAT**C**AATT TTAA-3’<br>Rev: 5’-CCTGTATTTTACCATACTTTGTTTTATTATTAAAATT**G** ATTTCTTTCATCCTTTC-3’ |

**Supplementary Table 1. Primer sequences used in this study.** All primers were purchased from Integrated DNA Technologies (IDT) or Elim Biopharm and prepared with standard desalting purification methods.

| Tiamulin µg/mL | Colony # | Mutations | 2nd Codon |
|---|---|---|---|
| 400 Enrichment Round 1 | 2 (CfrV6) | Promoter, I26M, E351Stop, 3'UTR-INS | AAU |
| | 3 | **N2K**, S39G, I326V, Q346H, E351Stop | **AAA** |
| | 4 | **N2K**, S39G, N57D, S348C, E351Stop, 3'UTR-INS | **AAA** |
| | 5 | **N2K**, S18R, E266D, E351Stop | **AAA** |
| | 6 (CfrV2) | **N2K**, S39G, E351Stop | **AAA** |
| 500 Enrichment Round 1 | 1 | **N2K**, I26M, S273R, E351Stop | **AAA** |
| | 2 | **N2K**, S39G, N347K, S348Stop | **AAA** |
| | 3 (CfrV3) | **N2K**, I26M, S39G, E351Stop, 3'UTR-INS | **AAA** |
| | 5 | **N2K**, S39G, K198N, Q346Stop | **AAA** |
| | 6 | **N2K**, S39G, E351Stop, 3'UTR-INS | **AAA** |
| | 7 | *N2I*, S39G, Q202H, M301K, E351Stop, 3'UTR-INS | *AUU* |
| 600 Enrichment Round 1 | 1 (CfrV1) | **N2K**, I26M, E351Stop, 3'UTR-INS | **AAA** |
| | 2 | *N2I*, I26M, N73H, E351Stop | *AUU* |
| | 4 | **N2K**, N20S, K35R, S39G, L68F, N238D, L265H, E351Stop | **AAA** |
| | 5 | *N2I*, S39G, Q349Stop, 3'UTR-INS | *AUU* |
| 700 Enrichment Round 1 | 1 (CfrV7) | Promoter, S39G, E351Stop, 3'UTR-INS | AAU |
| | 2 | N5K, S39G, I233L, E351Stop | AAU |
| | 4 | *N2I*, I26M, S39G, G308V, E351Stop | *AUU* |
| | 5 | S39G, L68F, G115R, K198R, E351Stop, 3'UTR-INS | AAC* |
| | 6 | S39G, L289M, E351Stop, 3'UTR-INS | AAC* |
| | 7 | *N2I*, S39G, E351Stop, 3'UTR-INS | *AUU* |
| | 8 | **N2K**, I26M, N238D, E351Stop | **AAA** |
| 800 Enrichment Round 1 | 1 | **N2K**, S39G, Q346R, Q349Stop, 3'UTR-INS | **AAA** |
| | 2 | *N2I*, S39G, I233L, P259H, Q349Stop, 3'UTR-INS | *AUU* |
| | 4 | N5I, K35R, S39G, E351Stop, 3'UTR-INS | AAC* |
| | 5 | Promoter, L68F, S348N, E351Stop, 3'UTR-INS | AAU |
| | 6 | Promoter, I26M, K45Q, L68F, E351Stop, 3'UTR-INS | AAU |
| 1000 Enrichment Round 2 | 1 | Promoter, *N2I*, D23E, I26M, A305T, Q349Stop, 3'UTR-INS | *AUU* |
| | 2 | **N2K**, S39G, Q346R, E351Stop, 3'UTR-INS | **AAA** |
| | 3 | **N2K**, I26M, T62A, E351Stop | **AAA** |
| | 4 | N5K, S39G, A305T, E351Stop | AAU |
| | 6 | **N2K**, S39G, N65S, Q349Stop, 3'UTR-INS | **AAA** |
| | 7 (CfrV5) | *N2I*, S39G, E351Stop, 3'UTR-INS | *AUU* |
| | 8 | **N2K**, I26M, Q349Stop, 3'UTR-INS | **AAA** |
| 1250 Enrichment Round 2 | 2 | S39G, L68F, G115R, K198R, E351Stop, 3'UTR-INS | AAC* |
| | 4 | Promoter, Y127F, D234G, E351Stop, 3'UTR-INS | AAC* |
| | 5 | Promoter, I26M, S39G, Q72K, S85T, E351Stop, 3'UTR-INS | AAU |
| | 6 | **N2K**, I26M, N65S, Q349Stop, 3'UTR-INS | **AAA** |
| | 7 (CfrV4) | **N2K**, I26M, L68F, E351Stop, 3'UTR-INS | **AAA** |
| | 8 | Promoter, N5K, S39G, S273N, S277R, K315E, E351Stop | AAU |
| 1500 Enrichment Round 2 | 1 | Promoter, **N2K**, R17S, N73H, E351Stop | **AAA** |
| | 2 | **N2K**, S39G, S196G, E270K, G308R, K315R, E351Stop, 3'UTR-INS | **AAA** |
| | 4 | Promoter, **N2K**, S39G, Q346H, S348I, 3'UTR-INS | **AAA** |
| | 5 | Promoter, I26M, Q36L, N347K, S348Stop, 3'UTR-INS | AAU |
| | 6 | Promoter, *N2I*, D23E, I26M, A305T, Q349Stop, 3'UTR-INS | *AUU* |
| | 7 | **N2K**, S39G, E351Stop | **AAA** |
| | 8 | Promoter, S39G, E351Stop, 3'UTR-INS | AAU |

**Supplementary Table 2. Evolved Cfr sequence variants observed during final enrichment rounds of directed evolution.** Open reading frame mutations and alterations to sequences 5' (promoter) and 3' (insertion in 3' untranslated region) of the *cfr* gene are designated. Green lettering designates to the original Asn codon in CfrWT, while green with * designates an Asn synonymous codon. N2K(AAA) codon is in red, while N2I(AUU) codon is in blue.

| Enrichment Round | Tiamulin µg/mL | Colony # | Promoter Architecture |
|---|---|---|---|
| - | - | CfrWT | Ptet – *cfr* |
| 1 | 400 | 2 (CfrV6) | Ptet – Ins – Ptet – *cfr* |
| | 700 | 1 (CfrV7) | Ptet – Ins – pPtet – *cfr* |
| | 800 | 5 | Ptet – Ins – pPtet – *cfr* |
| | 800 | 6 | Ptet – Ins – pPtet – *cfr* |
| 2 | 1000 | 1 | Ptet – Ins – pPtet – *cfr* |
| | 1250 | 4 | Ptet – Ins – pPtet – *cfr* |
| | 1250 | 5 | Ptet – Ins – pPtet – *cfr* |
| | 1250 | 8 | Ptet – Ins – pPtet – *cfr* |
| | 1500 | 1 | Ptet – Ins – pPtet – *cfr* |
| | 1500 | 4 | Ptet – Ins – pPtet – *cfr* |
| | 1500 | 5 | Ptet – Ins – pPtet – Ins – pPtet – *cfr* |
| | 1500 | 6 | Ptet – Ins – pPtet – *cfr* |
| | 1500 | 8 | Ptet – Ins – pPtet – *cfr* |

**Supplementary Table 3. Promoter architecture of evolved Cfr variants from final enrichment rounds with promoter alterations.** Abbreviations: Ptet = promoter; Ins = insertion sequence of various length; pPtet = partial promoter.

| MIC µg/mL | | | | | |
|---|---|---|---|---|---|
| **pZA** | **TIA** | **CHL** | **CLI** | **LZD** | **TMP** |
| Empty | 400-800 | 4-8 | 200 | 8 | 0.5-1 |
| CfrWT | 800-1600 | 8 | 400 | 16 | 0.5-1 |
| CfrV1 | 3200-6400 | 16 | 800 | 32 | 0.5-1 |
| CfrV2 | 3200-6400 | 16 | 800 | 32 | 0.5-1 |
| CfrV3 | 6400 | 16 | 1600 | 64 | 0.5-1 |
| CfrV4 | 3200-6400 | 16 | 800 | 32 | 0.5-1 |
| CfrV5 | 6400 | 16-32 | 3200 | 32 | 0.5 |
| CfrV6 | 1600-3200 | 16 | 800 | 32 | 0.5 |
| CfrV7 | 6400, >6400 | 64 | 6400 | 128 | 0.5 |

**Supplementary Table 4. Antibiotic susceptibility testing of Cfr variants.** Minimum inhibitory concentration (MIC) required to inhibit bacterial growth of *E. coli* BW25113 transformed with pZA plasmid encoding CfrWT or evolved Cfr variants. Antibiotic susceptibility testing was performed in biological triplicate by microbroth dilution method for tiamulin (TIA), chloramphenicol (CHL), clindamycin (CLI), linezolid (LZD), and trimethoprim (TMP). LZD testing was performed against *E. coli* BW25113 lacking efflux pump, *acrB*.

| pZA | TIA MIC µg/mL |
|---|---|
| Empty | 400-800 |
| CfrWT-GS1-FLAG | 1600 |
| CfrV1-GS1-FLAG | 6400 |
| CfrV2-GS1-FLAG | 6400 |
| CfrV3-GS1-FLAG | 3200-6400 |
| CfrV4-GS1-FLAG | 6400 |
| CfrV6-GS1-FLAG* | 3200 |
| CfrV7-GS1-FLAG* | 6400 |

**Supplementary Table 5. Antibiotic susceptibility testing of FLAG constructs.** Minimum inhibitory concentration (MIC) required to inhibit bacterial growth of *E. coli* BW25113 transformed with pZA plasmid encoding evolved Cfr variants with a C-terminal glycine-serine linker followed by a FLAG tag. Antibiotic susceptibility testing was performed in two biological replicates by microbroth dilution in the presence of tiamulin (TIA). Asterisk denotes that the 3' insertion sequence after the stop codon (3'UTR), which was introduced during directed evolution, was removed to install the C-terminal tag. Lack of the 3' UTR insertion sequence did not impact resistance for CfrV6/7.

| | |
|---|---|
| **Data collection and processing** | |
| Electron microscope | Krios |
| Magnification | 29 000 |
| Number of micrographs | 2055 |
| Number of particles picked from good micrographs | 162 713 |
| Number of particles used in final reconstruction | 141 549 |
| Pixel size (Å) | 0.822 |
| Defocus range (µm) | –0.2 to –1.5 |
| Voltage (kV) | 300 |
| Electron dose (e-/Å2) | 80 |
| **Map refinement** | |
| Model resolution (Å) | 2.2 |
| FSC threshold | 0.143 |
| Model resolution range (Å) | 2.2–20 |
| Map sharpening B-factor (Å2) | -55.86 |
| **Refinement and model statistics** | |
| Clashscore, all atoms | 2.23 |
| Protein geometry | |
| MolProbity score | 1.29 |
| Rotamer outliers (%) | 0.92 |
| Cβ deviations >0.25 Å (%) | 0.32 |
| Ramachandran (%) | |
| - Favored | 95.79 |
| - Allowed | 4.01 |
| - Outliers | 0.2 |
| Deviations from ideal geometry | |
| - Bonds (%) | 0.03 |
| - Angles (%) | 0.08 |
| Nucleic acid geometry | |
| Probably wrong sugar puckers (%) | 0.84 |
| Bad backbone conformations (%) | 12.86 |
| Bad bonds (%) | 0.07 |
| Bad angles (%) | 0.08 |

**Supplementary Table 6. Cryo-EM data collection, refinement, and validation statistics.**
